# Machine learning optimization of peptides for presentation by class II MHCs

**DOI:** 10.1101/2020.08.18.256081

**Authors:** Zheng Dai, Brooke D. Huisman, Haoyang Zeng, Brandon Carter, Siddhartha Jain, Michael E. Birnbaum, David K. Gifford

**Affiliations:** Computer Science and Artificial Intelligence Laboratory, MIT, Cambridge, MA, USA; Dept. of Computer Science and Electrical Engineering, Cambridge, MA, USA; Dept. of Biological Engineering, MIT, Cambridge, MA, USA

**Author notes:** These authors contributed equally to this work (Zheng Dai, Brooke D. Huisman).

## Abstract

T cells play a critical role in normal immune responses to pathogens and cancer and can be targeted to MHC-presented antigens via interventions such as peptide vaccines. Here, we present a machine learning method to optimize the presentation of peptides by class II MHCs by modifying the peptide’s anchor residues. Our method first learns a model of peptide affinity for a class II MHC using an ensemble of deep residual networks, and then uses the model to propose anchor residue changes to improve peptide affinity. We use a high throughput yeast display assay to show that anchor residue optimization successfully improved peptide binding.

## Introduction

Peptide vaccines are promising therapeutics for cancer and infectious disease that stimulate T cells to attack tumor or virally infected cells. T cells surveille peptides displayed on the cell surface by Major Histocompatibility Complexes (MHCs), or Human Leukocyte Antigens (HLAs) in humans, and T cell-mediated killing is initiated by recognition of a foreign peptide bound to an MHC. Specifically, CD8^+^ “cytotoxic” T cells recognize peptides presented by Class I MHCs, and CD4^+^ “helper” T cells recognize peptides presented by Class II MHCs [1]. Peptide vaccines serve to amplify a T cell response to cells displaying disease-associated peptides, and have proven successful clinically for patients with cancer after eliciting CD8^+^ and CD4^+^ T cell responses [2]. To formulate a peptide vaccine, its constituent peptides are computationally selected from a collection of disease associated peptides, with the criteria for vaccine inclusion including sequence differences from self peptides and the ability to be displayed by MHCs [2-4].

To improve the effectiveness of a peptide vaccine, the display of its component peptides can be optimized through anchor residue changes. It has been observed that peptide vaccines containing sequences with modified peptide anchor residues can improve the tumor cell killing response of the adaptive immune system [5]. For peptides presented by class I MHCs, not all modifications to the antigen sequences improve the recognition of tumor-displayed peptides by the immune system [6]. However, in contrast to class I MHCs which present peptides in an arched conformation within a closed peptide-binding groove, class II MHCs have open grooves in which presented peptides are displayed in an extended conformation, which results in peptides binding in a highly conserved manner. The peptide side chains at positions P1, P4, P6, and P9 are completely buried within binding pockets in the groove and are considered anchor positions [7]. These four anchor residues are key determinants of peptide-MHC binding affinity. Changing the identities of the class II MHC-binding anchor residues will allow us to alter binding affinity without changing binding conformation or T cell receptor contacts.

A rule based approach, EpitOpimizer, has been used to design modified peptides with anchor position changes that resulted in improved adaptive immune system response [8]. EpitOptimizer uses a limited sequence context for its suggestions, and each MHC Class I molecule has a different set of rules. By contrast, PeptX [9] uses a genetic algorithm to determine the peptides mostly likely to be displayed by a specific MHC class I allele, which may provide helpful information for the subsequent design of a vaccine. The performance of PeptX was not experimentally evaluated.

We introduce a model-based approach to optimize peptide-class II MHC binding by changing the peptide’s anchor residues. This approach uses the entire sequence of disease associated peptides to produce new peptide sequences with optimized MHC anchor residues. We adapt a yeast display platform for testing our improved peptide sequences for binding to class II MHC molecules. We optimize peptide-MHC class II affinity by enumerating all possible changes to the anchor positions of a peptide, then scoring them against an objective function in silico and choosing the best ones. This is computationally tractable due to the limited number of anchor positions on a given peptide.

For our objective function, we use predictions from the PUFFIN peptide-MHC binding model [10] trained on peptide binding data from a class II MHC yeast display platform [11]. PUFFIN uses an ensemble of deep residual networks that takes as input the peptide and MHC amino acid sequences and outputs a predicted affinity distribution of the peptide for the MHC, achieving state of the art performance on class II MHC binding prediction tasks [10]. We show that our method generates peptide modifications that improve peptide binding affinity for two class II MHCs.

## Results

### We evaluate the complete anchor substitution landscape with a machine learning model

For a given MHC class II molecule, we train a neural network-based machine learning (ML) model (PUFFIN) [10] that takes a 9 residue peptide sequence as input, and outputs a measure of the strength of the peptide-MHC interaction. PUFFIN outputs uncertainty estimates which allows us to compute Bayesian acquisition functions. We leverage the relatively small space of 20^4^-1 possible anchor substitutions to evaluate an objective function over each substitution based on the output of the model. We then output the 10 substitutions that score the highest as the proposed optimizations. The use of a neural network-based model along with the complete enumeration of the anchor substitution space allows our optimizations to take more complex interactions between residues into account.

### Data was collected using a high throughput peptide display assay that measures enrichment as a surrogate for affinity

We utilize peptide-MHC binding data from a yeast display platform [11] for data collection (Fig. 1). In this platform, class II MHCs are covalently linked to a query peptide with a flexible linker which contains a 3C protease cleavage site. When the linker is cleaved, unbound peptides can be displaced from the MHC in the presence of a high-affinity competitor peptide. The linker also contains a peptide-proximal epitope tag, which we use to enrich yeast that maintain peptide-MHC binding. Data is collected over multiple iterative rounds of selection. After each round of selection, deep sequencing is carried out on the enriched sequences.

**Figure 1.**
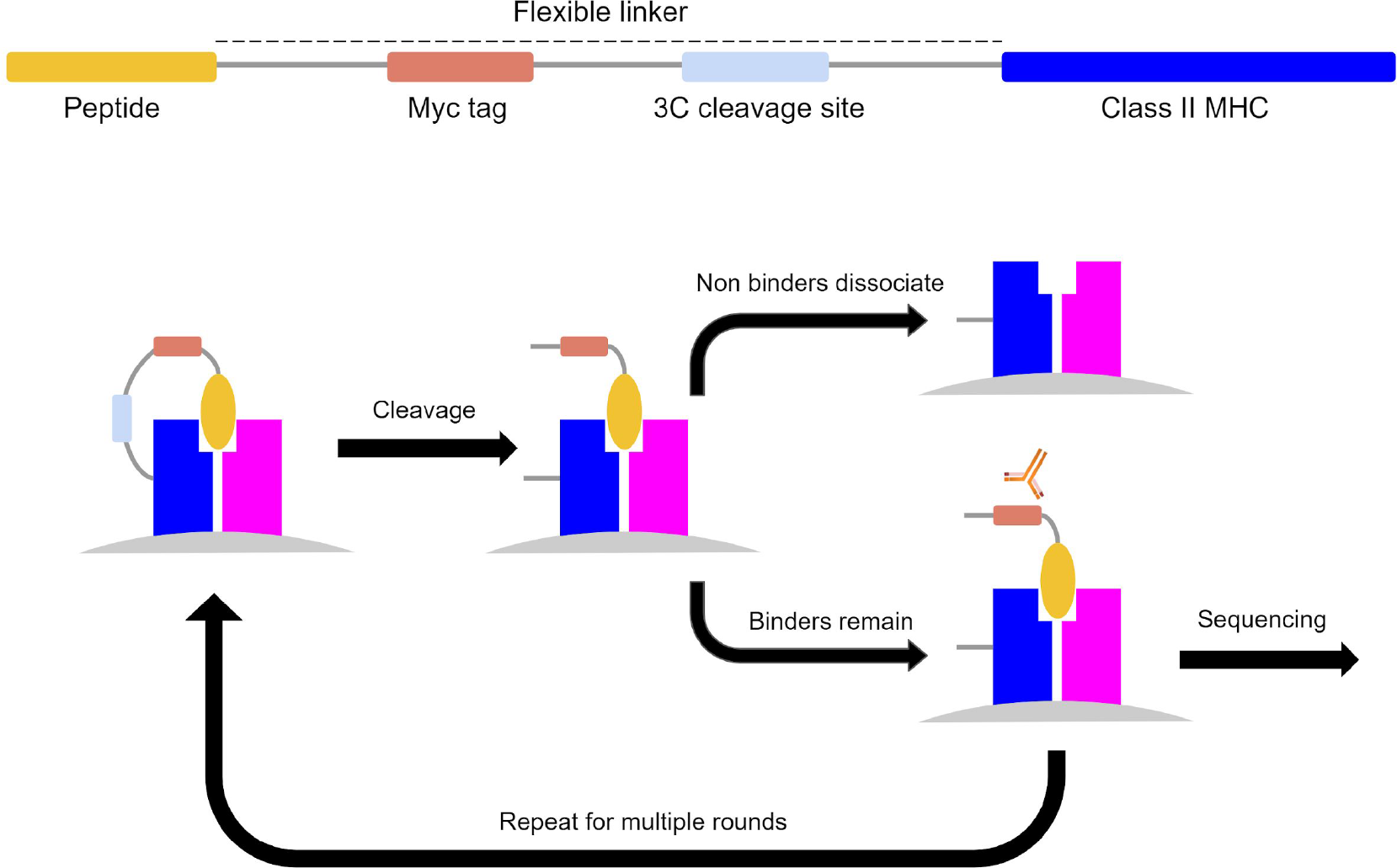
Characterization of peptide class II MHC binding by yeast display. On the top is a schematic of the construct used in the assay. The diagram on the bottom shows the overall process. First, the peptide-MHC is expressed on the surface of yeast, and then the linker between peptide and the MHC molecule is cleaved. Peptide exchange is catalyzed, and yeast are selected which retain the Myc epitope tag. The resulting population is then sequenced and carried on to the next round.

We utilized data from two class II MHC alleles: HLA-DR401 (HLA-DRA1*01:01, HLA-DRB1*04:01) and HLA-DR402 (HLA-DRA1*01:01, HLA-DRB1*04:02). This ensures that our results are not an allele specific artifact and allows us to study optimization for multiple alleles.

We utilized enrichment data from a library consisting of random 9-mer peptides flanked by invariant peptide flanking residues (IPFR) which encourages binding in a single register and simplifies identification of anchor residues [11]. We used this data to train 2 predictors for each allele. The first predictor models the enrichment as a continuous value and outputs a Gaussian distribution, while the second predictor models the enrichment as categoricals and outputs a probability distribution over the categories. In both cases, the enrichment value of a given 9-mer is based on the last round the 9-mer appears in.

### We optimized the anchor residues of sequences drawn from viral proteomes

We proposed anchor optimizations to 9-mers drawn from the proteomes of the Zika, HIV, and Dengue viral proteomes, which we refer to as seed sequences. We selected three sets of sequences on which to evaluate three different optimization tasks:

1. 82 seed sequences that originally bind mildly to HLA-DR401 were optimized for high affinity to HLA-DR401.
2. 87 seed sequences that originally bind mildly to HLA-DR402 were optimized for high affinity to HLA-DR402.
3. 44 seed sequences that originally bind strongly to HLA-DR402 but mildly to HLA-DR401 were optimized for high affinity to both MHC alleles.

PUFFIN was designed to characterize the uncertainty of its predictions by outputting a variance. This allows us to use various Bayesian acquisition functions as our objectives. For this study, we chose to study point estimate (PE) which is just the enrichment, and upper confidence bound (UCB) which adds the enrichment and the standard deviation of the prediction. For our third task of optimizing for both alleles, the objectives were computed for each allele individually and then added to produce the combined objective.

Optimizations using PE and UCB were performed with both the Gaussian and categorical models, giving a total of 40 optimized sequences for each seed. The optimized sequences and their seed sequences were then flanked with IPFR and added to a new yeast display library for testing our designs. For each seed, 10 random anchor substitutions were also generated as a random control and flanked with IPFR. As a further control, we also flanked all the 9-mers with their wild type peptide flanking residues (WPFR), which were defined as the 3 residues that flanked the seed 9-mer in the source proteome. Finally, we sampled some sequences that performed well and some sequences that performed poorly in the training data and added them as positive and negative controls respectively.

We constructed a new yeast display library composed of these optimized peptides and controls, and we conducted another series of selections to enrich for binders.. To compare affinities between given peptides, for each peptide we estimated the proportion of that peptide which survives between rounds. This proportion is unnormalized, so we refer to it as a round survival rate (RSR). We use it as a surrogate for affinity for the remainder of this section.

### Optimization improves the round survival rate of peptides for both HLA-DR401 and HLA-DR402

We first examine the RSR distribution of the following groups of sequences for each allele: sequences optimized for that allele with PE under the Gaussian model, UCB under the Gaussian model, PE under the categorical model, UCB under the categorical model, sequences with random anchor mutations (negative control), seed sequences (negative control), sequences from the training data which were not present after round 2 (negative control), and sequences from the training data which were present after round 2 (positive control). We find that the groups of optimized sequences exhibit higher RSRs for the alleles they were optimized for than either of the negative controls (Fig. 2). The improvements are statistically significant, with p ≤ 1.58e-23 between any optimized set and negative control for either allele by the two sided Mann-Whitney U test.

**Figure 2.**
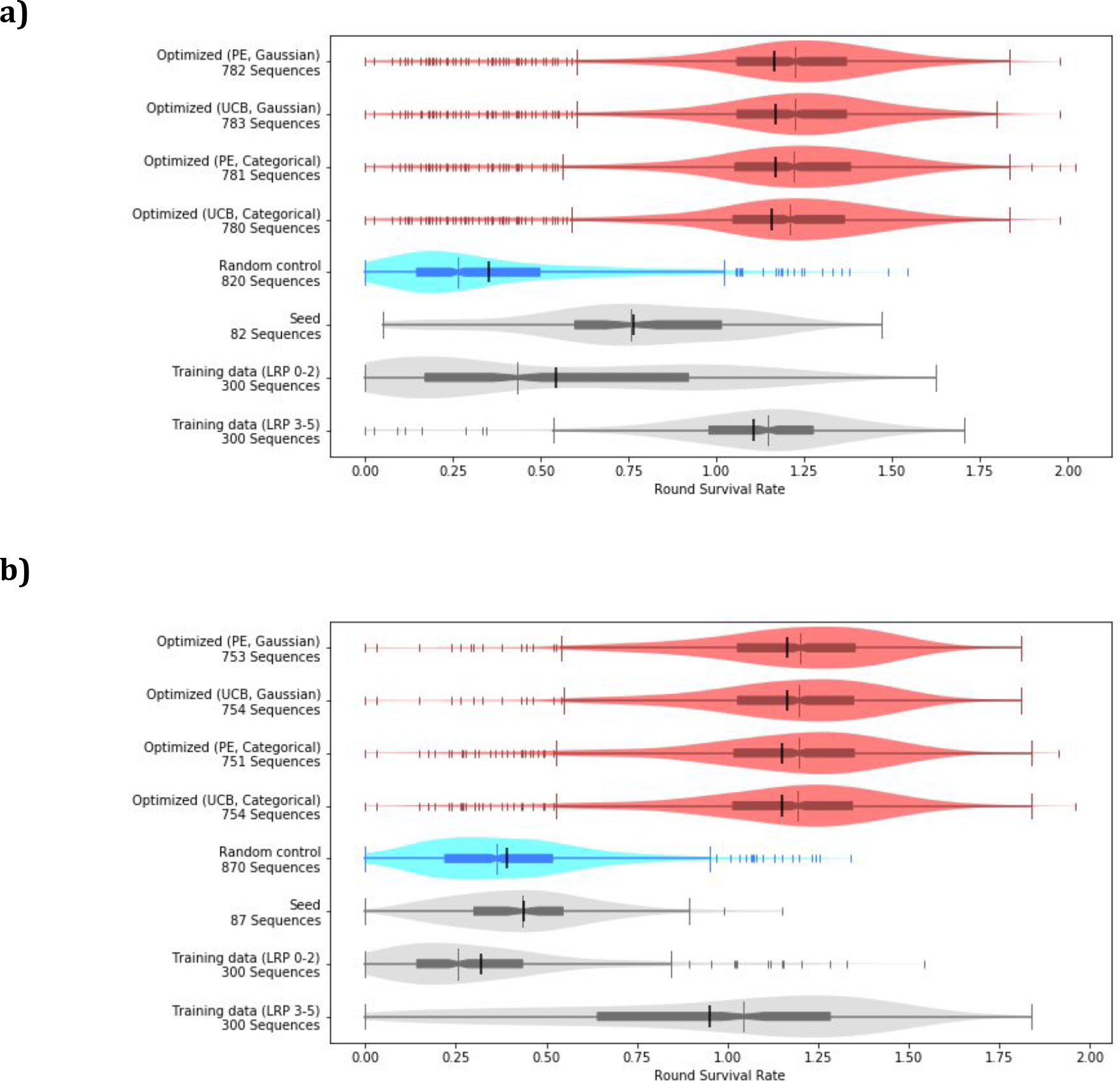
Anchor optimization improves round survival rate. The distributions of RSR for **a)** HLA-DR401 and **b)** HLA-DR402 is plotted for the optimized and control groups. The sequences from the training data are split into two groups: “Training data (LRP 0-2)” is composed of sequences which did not appear after round 2 in the initial display assay and is shown as a negative control, while “Training data (LRP 3-5)” is composed of sequences that did appear after round 2 and is shown as a positive control. The differences between the optimized groups and the negative controls (Random control, Seed and Training data (LRP 0-2)) are significant for both alleles, with p ≤ 1.58e-23 under the two sided Mann-Whitney U test. Each plot is a combination of a box plot and a violin plot, where the distribution is shown by the violin plot in a lighter color, and the box plot shows the middle quartiles in a darker color along with the median. The mean is indicated by a black vertical line. Flier points are marked with the “|” symbol.

In aggregate, the optimized sequences outperform the unoptimized sequences with comparable results for all of our optimization methods. For simplicity, we will focus the rest of this section on our point estimate optimization under the Gaussian model. We include analysis of the other optimization methods, which are similar, in the supplemental section (Supplemental Figures 1-4).

We find that most optimized sequences outperform their unaltered seed sequences (Fig. 3a, c). For HLA-DR401, for 44 out of the 82 seed sequences, all of the proposed optimizations performed better, while for HLA-DR402 this was the case for 72 out of the 87 seed sequences. In sequences where optimization was less effective, we find that generally the seed sequence already has a decent round survival rate (Fig. 3b, d).

**Figure 3.**
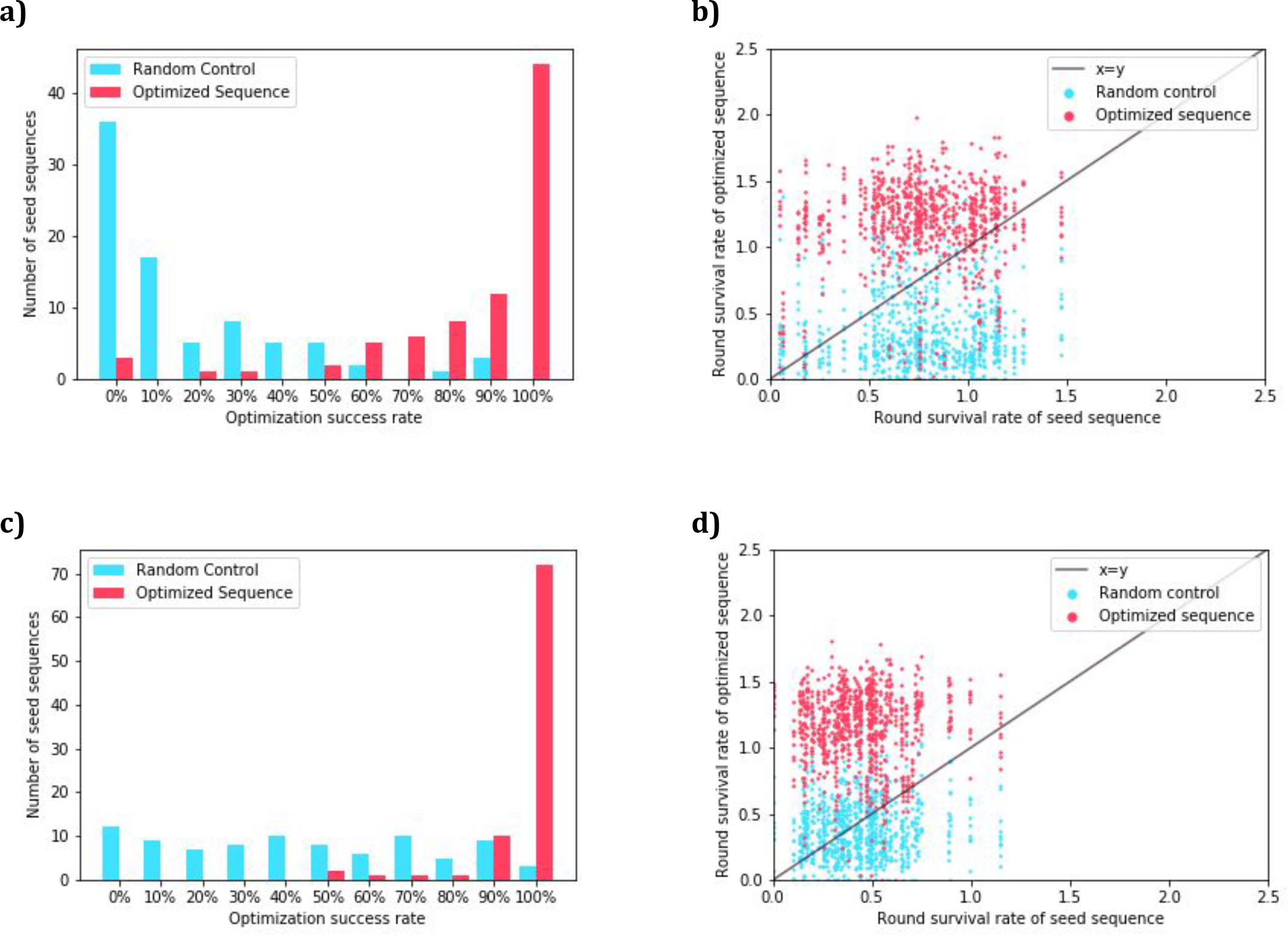
Number of sequences that exhibit improvement after optimizing with the point estimate objective under the Gaussian model. **a)** For each seed sequence, we calculate for each seed sequence the number of proposed optimizations that achieve a RSR for HLA-DR401 that is higher than that of the seed. We then take that as a percentage of the number of proposals to obtain the optimization success rate. We plot the distribution of these rates for both sequences optimized for HLA-DR401 affinity and the randomly perturbed sequences. **b)** For each sequence optimized for HLA-DR401 affinity and randomly perturbed sequence, we plot their RSR for HLA-DR401 against the RSR of the seed sequence they derive from. **c)** We calculate the distribution of optimization success rates for sequences optimized for HLA-DR402 using RSR for HLA-DR402. **d)** We plot the RSR for HLA-DR402 of sequences optimized for HLA-DR402 against the RSR of their seed sequence.

### Sequences can be optimized for multiple alleles simultaneously

We find that sequences that were optimized for both alleles were generally able to improve their RSR for HLA-DR401 while maintaining their RSR for HLA-DR402 (Fig 4). Out of a total of 44 seed sequences, there were 35 in which all proposed optimizations had a higher RSR for HLA-DR401. For 23 seed sequences, all proposed sequence optimizations outperformed the seeds on HLA-DR401 and achieved greater than 80% of the seed sequence RSR for HLA-DR402. For 13 seed sequences, all proposed optimizations outperformed the seeds on both HLA-DR401 and HLA-DR402.

**Figure 4.**
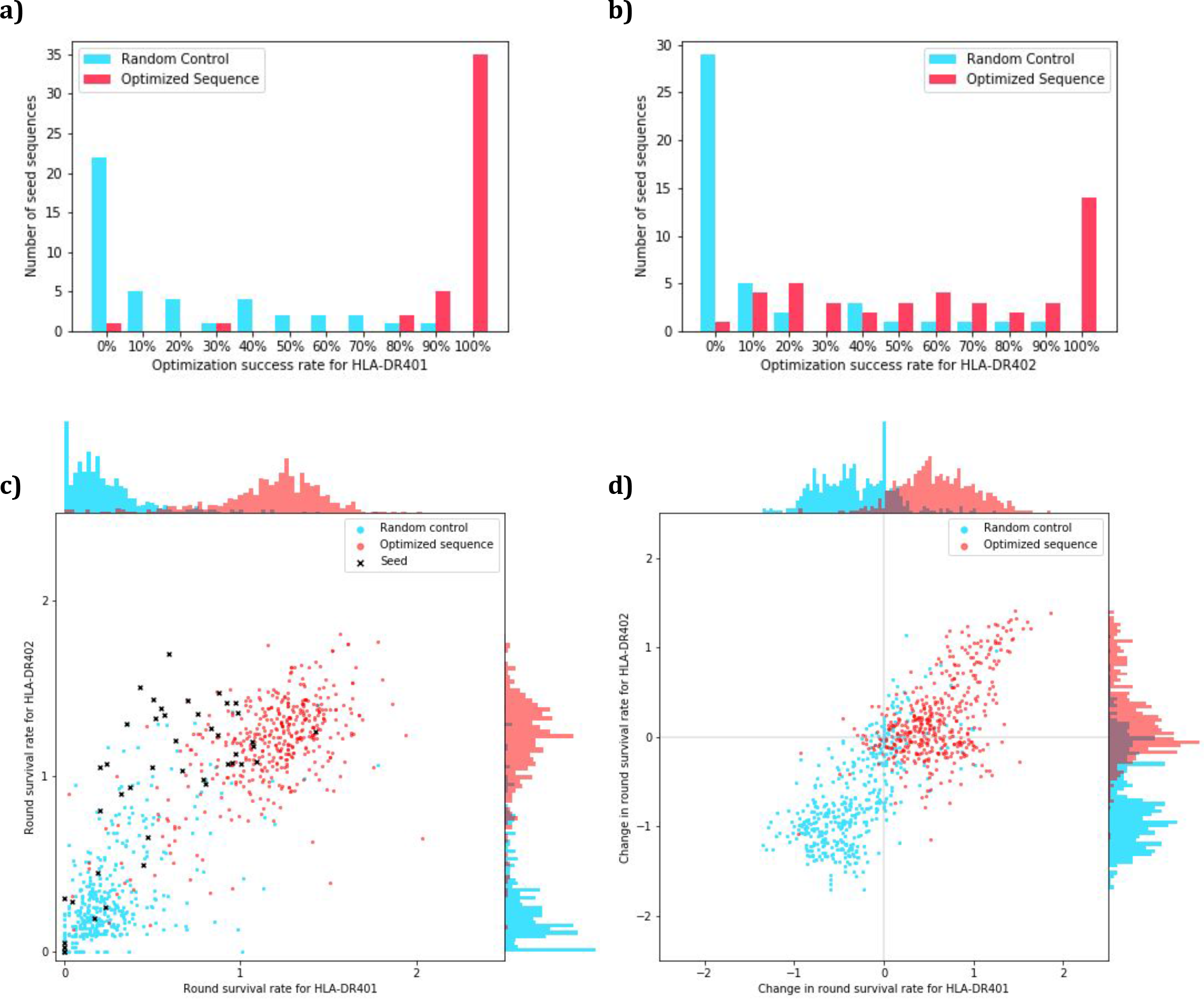
Number of sequences that exhibit improvement for multiple alleles after optimizing with the point estimate objective under the Gaussian model. **a)** For each seed sequence, we calculate for each seed sequence the number of proposed optimizations that achieve a RSR for HLA-DR401 that is higher than that of the seed. We then take that as a percentage of the number of proposals to obtain the optimization success rate. We plot the distribution of these rates for both sequences optimized for HLA-DR401 and HLA-DR402 affinity and the randomly perturbed sequences. **b)** We produce the same distribution but with optimization success rates based on HLA-DR402 affinity. The seed sequences were selected to have high HLA-DR402 affinity. **c)** For each optimized, random control, and seed sequence, we plot their RSR for both alleles. **d)** For each optimized and random control sequence, we take their RSR and subtract the RSR of the seed sequence they derive from to obtain the changes in their RSR.

If we instead consider seeds where the optimization criterion was reached for at least 8 out of the 10 proposed sequences, these values rise to 42, 35, and 18 respectively. For the random controls, they are 2, 1, and 1 (Fig. 4).

### Our optimizations capture complex interactions between residue positions

By analyzing our training data, we find that the identity of residues outside of the primary anchor residues can have a significant impact on which anchor residues will improve affinity. As an example (Fig. 5), for HLA-DR401 if a sequence contains a threonine (T) at the non-anchor position P7, then having an aspartic acid (D) at anchor position P6 tends to increase the RSR. However, if the sequence contains a D at the non-anchor position P7, then having a D at P6 tends to decrease the RSR instead. Higher order effects can be seen between other anchor and non-anchor positions as well, so these relationships are not limited to adjacent positions nor to residues at P7, which can be considered an auxiliary anchor because of its contacts with the MHC groove [7].

**Figure 5.**
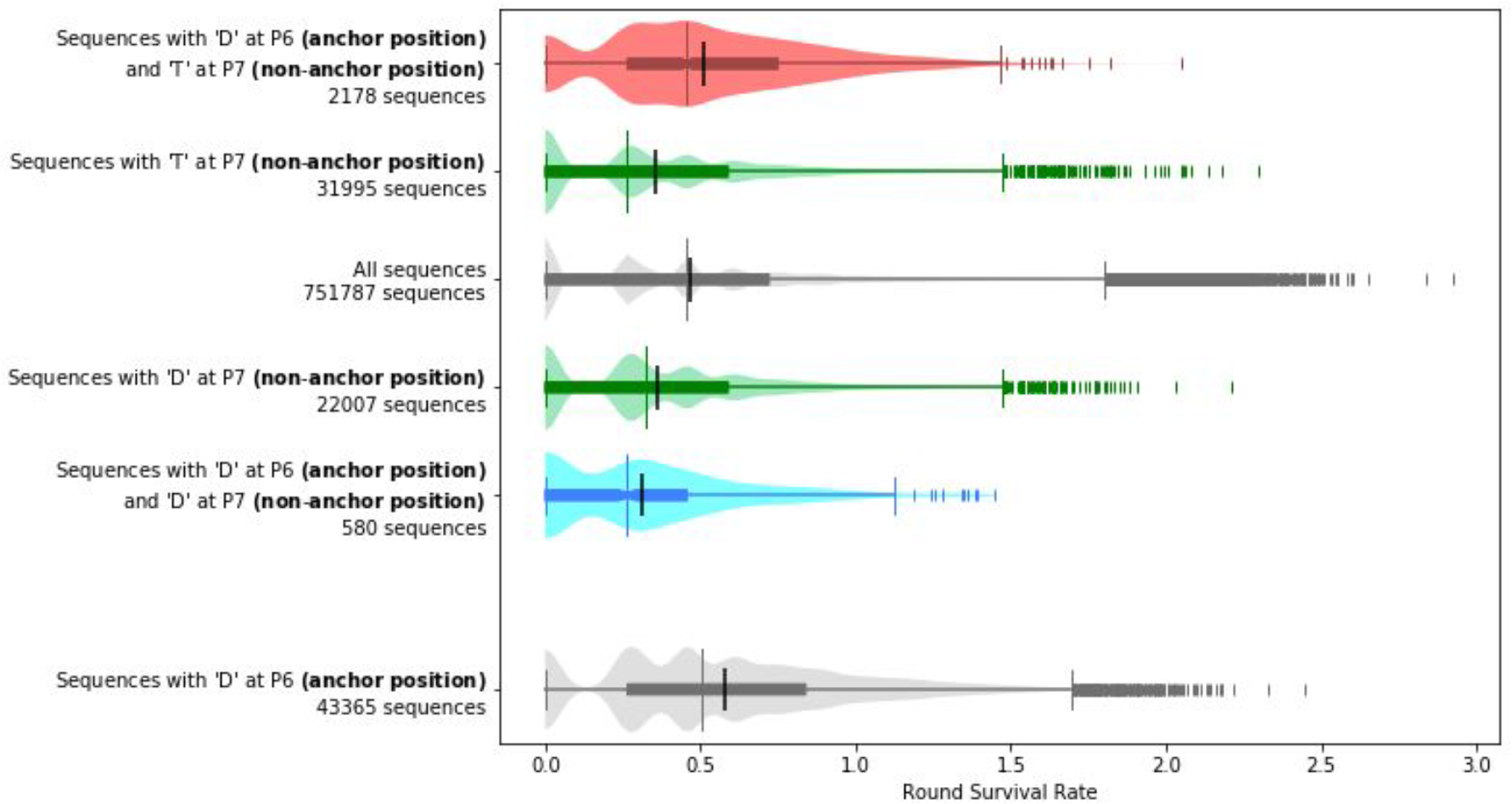
Whether a given anchor residue improves affinity can depend on non-anchor residues. Peptides with aspartic acid at P7 tend to have lower round survival rates when compared to all peptides, and peptides that additionally have another aspartic acid at anchor position P6 tend to have even lower round survival rates than the peptides that just have an aspartic acid at P7. In contrast, although peptides with threonine at non-anchor position P7 also tend to have lower round survival rates when compared to all peptides, peptides that additionally have an aspartic acid at anchor position P6 tend to have higher round survival rates instead even when compared to all peptides. The differences found in these comparisons are significant, with p ≤ 4.968e-5 between any two groups mentioned above under the Mann-Whitney two sided U test. The sequences plotted and used for computing significance are from the training data. Each plot is a combination of a box plot and a violin plot, where the distribution is shown by the violin plot in a lighter color, and the box plot shows the middle quartiles in a darker color along with the median. The mean is indicated by a black vertical line. Flier points are marked with the “|” symbol.

The dependency between anchor positions and non-anchor positions can be observed in the proposals generated by our method. Out of the 820 sequences proposed using PE under the Gaussian model for HLA-DR401, 20 (2%) have a D at anchor position P6. Six of our seed sequences have a T at P7; of our proposed optimizations for these seeds, 17/60 (28%) have a D at P6. Conversely, nine of our seed sequences have a D at P7, and none of their 90 proposed optimizations have a D at P6. This demonstrates the advantages of enumerating the full anchor residue landscape as it allows the capture of these higher order effects. In general, the predicted enrichments for each peptide that PUFFIN generates correlates strongly with the measured RSR (Supplemental Figure 5).

Given the presence of the higher order effects between peptide positions, including non-anchor positions, it seems unlikely that a more naive approach to anchor optimization could be as successful. In particular, it is unlikely that there exists a set of anchor residues that would optimize affinity in all non-anchor contexts. As further support for this, we find that there are no sets of anchor residues that were proposed for all seed sequences for any optimization task, even when combining the proposed optimizations across all 4 of our optimization methods. For HLA-DR401 optimization, the most frequently proposed set is Y, D, T, A at anchor positions P1, P4, P6, P9 (respectively), which was proposed for 54 out of the 82 seeds. For HLA-DR402, the most frequently proposed set is L, W, T, A at P1, P4, P6, P9 (respectively), which was proposed for 44 out of the 87 seeds. For optimization for both alleles, the most frequently proposed set is F, M, N, A at P1, P4, P6, P9 (respectively), which was proposed 34 out of 44 times.

### Optimized sequences outperform seed sequences in the absence of the invariant flanking residues, but is less effective

All the optimized peptides we have presented so far have been flanked with IPFR, which were used to train the model. If we replace the IPFR with WPFR, we observe that the optimized sequences still outperform the seed and random controls (Supplemental Figure 6). The improvement is still significant, with p ≤ 5.51e-5 when comparing the optimized sequences to the random control or seed sequences for either allele under the two sided Mann-Whitney U test. However, the optimized sequences with WPFR significantly underperform their IPFR counterparts (p ≤ 1.36e-22 for either alleles under the two sided U test).

**Figure 6.**
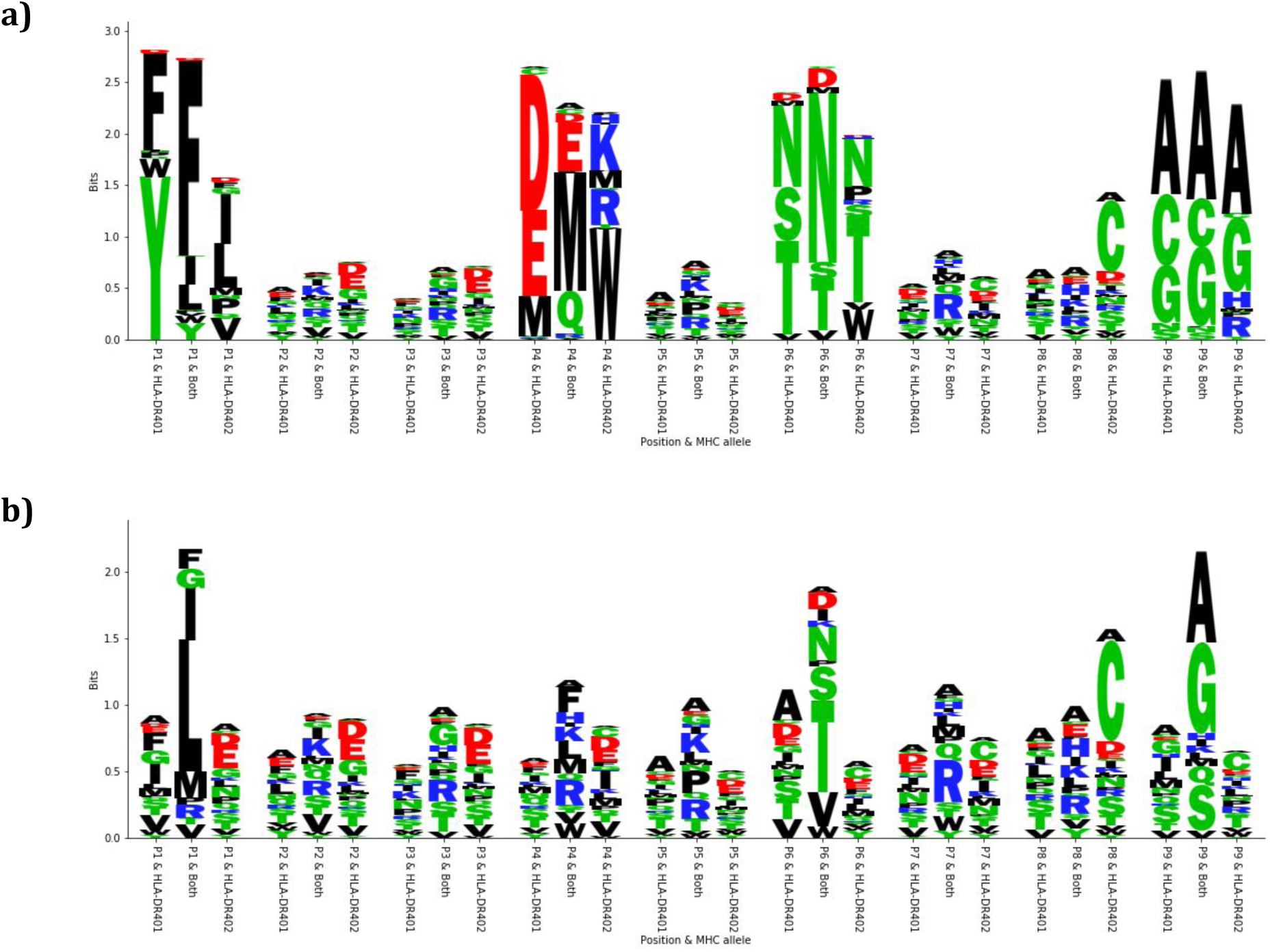
Motifs arising from optimization with the PE objective under the Gaussian model. **a)** Sequence logos depicting the motifs present in the optimized sequences. For each position, 3 different residue distributions are shown. The first one shows the distribution for the sequences optimized for HLA-DR401, the last one shows the distribution for the sequences optimized for HLA-DR402, and the one in the middle shows the distribution for the sequences optimized for both alleles. **b)** Sequence logos depicting the motifs present in the original seed sequences for comparison with the same setup.

In the case of the seed and random control groups, the WPFR sequences either do not display any significant difference or mildly outperform the IPFR counterparts (0.0018 ≤ p ≤ 0.92 under the two-sided Mann-Whitney U test). The drop in RSR is only observed in the optimized sequences.

### Optimized sequence motifs are consistent with MHC binding preferences

The peptide optimizations made by our machine learning models are consistent with the structures and peptide-binding motifs of HLA-DR401 and HLA-DR402 in our training data. The polymorphisms between HLA-DR401 and HLA-DR402 affect the P1 and P4 binding pockets. Both alleles prefer hydrophobic amino acids in the P1 pocket, although HLA-DR401 prefers larger amino acids, while the truncated HLA-DR402 pocket prefers smaller amino acids. In the P4 pocket, HLA-DR401 prefers acidic residues, and HLA-DR402 prefers basic residues and large hydrophobic residues. The conserved P6 and P9 binding pockets prefer polar and small amino acids, respectively. The preference for each allele is reflected in MHC allele-specific peptide optimization, shown for the optimization with PE objective under the Gaussian model as an example (Fig. 6a). Joint MHC optimization is also consistent with these preferences: P1 and P4 amino acids are mutually preferred between both alleles, such as F/I/L at P1 and increased usage of M at P4. P6 and P9 amino acids are consistent with usage in individual allele-optimized peptides. Amino acid frequency in the seed sequences are also shown for reference (Fig. 6b).

## Discussion

In this work, we introduced our method for optimizing the affinity of peptide sequences for class II MHCs by replacing their anchor residues with more optimal residues generated with the help of a machine learning model. We validated this technique on two different class II alleles, and showed that it is possible to optimize a single sequence for multiple alleles simultaneously. We have developed a high throughput yeast display-based pipeline to test our optimized sequences, and we introduced the notion of a round survival rate which allows us to compare the results of the assay.

We demonstrated that our method leverages a deep learning model in a way that allows our optimizations to capture complex interactions between residues. Our ability to optimize sequences for two alleles simultaneously suggests that the method can generalize to even more complicated objectives. These contributions improve our ability to engineer peptides for therapeutic purposes, and allows us to develop more robust cocktails by allowing their constituent peptides to fulfill multiple objectives.

As a caveat, we note that our optimization is less effective if we allow arbitrary flanking residues. The drop in RSR when we change from IPFR to WPFR is only observed in the optimized sequences and is not observed in seed or sequences with random anchor residues. Therefore, it is likely that the drop in performance is due to the predictor being trained on IPFR data, so the predictor is unable to take the effects of flanking residues or register shifts into account. The IPFRs also contain preferred amino acids in the flanking sequences, such as the tryptophan at position P10. Aromatic residues at P10 have been shown to bolster binding and may impact the superior performance of IPFR peptides compared to WPRF peptides [11, 12]. We note that since our method is independent of the specific underlying predictor, we should be able to address this issue by replacing our current predictor with one that takes the flanking residues into account. As a general statement, the quality of our optimization should improve as the quality of predictors available continues to improve.

Our future work will attempt to address the issues pertaining to the sophistication of the predictor, and will seek to characterize the effect of anchor optimization on peptide immunogenicity.

## Methods

### Collecting enrichment data using a high throughput peptide display assay

We utilize peptide-MHC binding data from a yeast display library of ∼ 10^8^ random 9mer peptides (Fig. 1)[11]. The peptides are flanked by invariant peptide flanking residues (IPFR), which encourages binding in a single register and simplifies identification of anchor residues. The IPFR consists of “AA” on the N-terminus and “WEEG” on the C-terminus. Paired-end sequencing reads [11] were assembled via FLASH [13] and filtered for correct length and 3C cut site sequence.

In order to test our optimized sequences, we adapted the yeast display platform and workflow from randomized peptide libraries to presentation of user-defined peptides. We designed a 36,000-member defined library containing our optimized sequences, which was synthesized by Twist Bioscience as a single-stranded oligo pool with a maximum length of 120 nucleotides. The oligo pool was amplified with low cycle number PCR then amplified with construct DNA using overlap extension PCR. This longer DNA product was assembled with the linearized pYal vector in yeast at a 5:1 mass ratio of insert:vector and electroporated into electrocompetent RJY100 yeast. To better assess enrichment, the HLA-DR401 and HLA-DR402 defined peptide libraries were doped into a ∼ 20-million-member randomized peptide library containing stop codons at a ratio of approximately 1:500 so that each unique peptide was represented at similar starting frequency. The diverse null library had the peptide encoded as “NNNTAANNNNNNNNNTAGNNNNNNNNNNNNTGANNNNNN”, where N indicates any nucleotide. Doping into this library provides a null set of peptides over which real binders must enrich.

For each round of selection, yeast were washed into PBS, with competitor peptide (HLA-DR401: HA_306-318_, 1uM; HLA-DR402: CD48_36-53_, 5uM) and 1uM 3C protease, then incubated for 45min room temperature. After incubation, yeast were washed into cold acid saline (20mM pH5 citric acid, 150mM NaCl) with competitor peptide (same concentration as first incubation) and 1uM HLA-DM, then incubated overnight at 4°C. Negative selections for non-specific binders was performed with anti-AlexaFluor647 magnetic beads (Milltenyi Biotech; Bergish Gladbach, Germany), followed by a positive selection consisting of incubation with anti-Myc-AlexaFluor647 antibody (1:100 volume:volume) and positive selection with anti-AlexaFluor647 magnetic beads. The first round was conducted on 400 million yeast for 20x coverage of peptides and incubations were conducted in 2mL PBS and 4mL acid saline. For subsequent rounds, 25 million yeast were selected; incubations were conducted in 250uL PBS, 500uL acid saline. Four iterative rounds of selection were performed and repeated in duplicate. Between rounds, yeast were grown to confluence at 30°C in SDCAA (pH=5) yeast media and subcultured into SGCAA (pH=5) media at OD_600_=1 for two days at 20°C [14].

Following selections, plasmid DNA was isolated from 10 million yeast from each round using a Zymoprep Yeast Miniprep Kit (Zymo Research; Irvine, CA). Amplicons were generated to capture the peptide through the 3C protease site. Unique barcodes were added for each library and round of selection and i5 and i7 anchors added through two rounds of PCR. Amplicons were sequenced on an Illumina MiSeq (Illumina; San Diego, California) at the MIT BioMicroCenter, with a paired-end MiSeq v2 300nt kit.

Forward and reverse reads are assembled using PandaSeq. Data were processed using in-house scripts to extract peptide sequences with correctly encoded constant flanking regions. Peptides were filtered for exact matches to the defined sequences ordered from Twist and those matching the DNA encoding of the randomized null library.

HLA-DM was recombinantly expressed as previously described [11]. In brief, the ectodomains of the alpha and beta chains were followed by a poly-histidine purification site and encoded in pAcGP67a vectors. Plasmids for each chain were separately transfected into SF9 insect cells with BestBac 2.0 baculovirus DNA (Expression Systems; Davis, CA) and Cellfectin II reagent (Thermo Fisher; Waltham, MA). Cells were propagated to high virus titer, co-titrated to ensure an equal ratio of alpha and beta expression, and co-transduced into Hi5 cells. Following 48-72 hours of incubation, proteins were purified with Ni-NTA resin and purified with size exclusion chromatography on an AKTAPURE FPLC S200 increase column (GE Healthcare; Chicago, IL).

### Training a neural network based ML model to predict the enrichment category of a peptide

We trained a neural network-based machine learning (ML) model (PUFFIN) [10] to predict the enrichment label of a new peptide. The predictor takes a 9 residue peptide sequence as input, and outputs an enrichment label. We use the final round where a peptide is observed in the peptide display assay as our enrichment label for both training and prediction. For example, if a sequence appears in the sequencing reads for round 3 but fails to appear in round 4 or any other future rounds, it receives the label “3”. To improve the granularity, round 5 presence was further split up into 3 categories, where “5” indicates round 5 presence with less than 10 read counts in round 5, “6” indicates round 5 presence with a read count between 10-99 inclusive, and “7” indicates round 5 presence with a read count of 100 or more. A label of 0 is given to sequences that only appear before any enrichment is performed. This gives a total of 8 enrichment categories.

PUFFIN is an ensemble of deep residual neural networks that is regularized by dropout and controlled for overfitting with validation data. Each component model consists of one convolutional layer, five residual blocks, and one output layer. Each residual fits the difference between the input and the output of a residual block with two convolutional layers. Each convolutional layer has 256 convolutional filters and is followed by a batch-norm layer. ReLU is used as non-linearity throughout the network.

For each allele, we trained two predictors to predict the enrichment labels. The first predictor assumes the enrichment labels 0-7 are realizations of a continuous random variable taken from a Gaussian distribution, and was trained to output a mean and variance. The second predictor models the labels as categoricals, and outputs a discrete probability distribution over the 8 labels 0-7. For regularization, dropout [15] is used in the output layer with a dropout probability of 0.2. We randomly hold out 10% of the data for validation, and the rest is used for training. We use Adam [16] to minimize the negative log-likelihood of the observed enrichment under the probability distribution parameterized by the output of the neural network. We train for 50 epochs and select the model from the epoch where validation loss is minimized.

While the outputs of each predictor naturally characterize aleatoric uncertainty, we also characterize the epistemic uncertainty through ensemble methods [17]. Specifically, we generate 10 training and validation splits of our data and train 2 separate predictors for each split, giving us an ensemble of 20 predictors. When performing predictions, we run each predictor 50 times with dropout turned on [18], resulting in a total of 1000 predictions for each input. The final output is then characterized by a mean and variance, where the mean is the average of the distribution means over all 1000 trials, and the final variance is the average of the distribution variances for each trial plus the variance of the distribution means.

### Using an ML model to compute an objective function for anchor optimization

For both the Gaussian and the categorical predictors, we considered two different objective functions for scoring 9-mers: point estimate (PE) and upper confidence bound (UCB). In both functions, we first run our predictor over the 9-mer to obtain a predicted mean and variance. Then to compute the PE objective, we simply return the mean. To compute UCB, we return the sum of the mean and the standard deviation, which we take to be the square root of the variance.

This gives us a total of 4 methods for scoring 9-mers. For each method, given an input 9-mer to optimize we enumerate all possible residue substitutions at positions 1, 4, 6, and 9 (sequences are 1-indexed). For each substitution, we compute its score using our objective function, and in the end we output the 10 sequences that score the highest as proposed optimizations.

### Designing a validation library to test the efficacy of anchor optimization

We evaluate our optimization methods on three tasks:

1. Optimizing peptide sequences that originally bind mildly to HLA-DR401 for higher affinity to HLA-DR401;
2. Optimizing peptide sequences that originally bind mildly to HLA-DR402 for higher affinity to HLA-DR402;
3. Optimizing peptide sequences that originally bind strongly to HLA-DR402 but mildly to HLA-DR401 for high affinity to both MHC alleles.

Our evaluation was conducted using viral peptides selected from the Zika, HIV, and Dengue viral proteomes. The 9-mers in the candidate proteomes have no overlap with the peptides in our random peptide training library. We selected seeds for optimization from Zika, HIV, and Dengue based on the predictions of the categorical predictor. We first filter the sequences by removing all whose PUFFIN prediction has a predicted variance higher than the median predicted variance. For the seeds for Task 1 and Task 2, we selected peptides with a predicted enrichment mean between 2 and 3, yielding 82 seeds for HLA-DR401 and 87 seeds for HLA-DR402. For the seeds for Task 3, we selected peptides with a predicted HLA-DR401 enrichment mean below 3 and a predicted HLA-DR402 enrichment mean above 5, resulting in 44 seeds.

For each seed sequence, we ran each of our 4 optimization methods over it for each allele, giving 10 optimized sequences for each method. As a control, we also proposed 10 random anchor residue mutations for each seed.

We then take the seed, optimized, and random sequences and flank them with IPFR. As a control, we also produce a second set of sequences from the same 9-mers but flanked with WPFR, defined to be the 3 residues that flank the original seed sequence in the original proteome. This forms the basis of the library.

As a further control, we added sequences from the original training data to the library. For each allele, we sampled 300 sequences that had no presence after the second round of selections, and 300 sequences that had presence after the second round of selections, giving us 1200 sequences overall.

### Calculating round survival rate as a representation of enrichment

The enrichment information reported in the peptide display assay comes in the form of a vector of read counts indexed by round. In order to compare enrichment between different peptides, we assign to each peptide a value that can be interpreted as an unnormalized proportion of that peptide that survives between rounds of enrichment. We will refer to this quantity as a round survival rate (RSR), where a higher RSR will be indicative of higher enrichment.

To calculate a peptide’s RSR, we consider a simplified model where the peptide has a starting concentration drawn from a given prior, and the dominating event is peptide dissociation from the class II MHC. Additionally, we assume that we can treat the entire experiment as though it was happening in one solution, and everything that occurs between rounds can be captured by a scaling factor. Finally, we suppose that read counts follow a Poisson distribution parameterized by the concentration multiplied by a scaling factor.

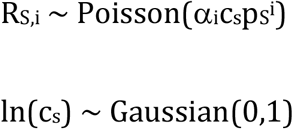

For all peptides S and all rounds i, where R_S,i_ is the read count of peptide S in round i, c_S_ is the starting concentration of the peptide S, p_S_ is an unnormalized proportion of peptide S that survive to the next round, and α_i_ is a round specific constant. The prior for constraining c_S_ is for regularization purposes, and a log normal distribution was selected for its interpretability as the outcome of a Wiener process.

We then define the RSR for peptide S as the maximum a posteriori (MAP) estimate of p_S_. This value is not unique, as an adequate scaling in the α_i_ values can give the same probabilities with different p_S_ values. However, such a transformation preserves the ratio between p_S_, and in practice we find that the estimates converge reliably (Supplemental Figure 7). We estimate these values by iteratively optimizing each variable individually for 500 rounds. 23 rounds were carried out with random initializations, where p_S_ were drawn from Uniform(0.1,1) and ln(c_S_) were drawn from Gaussian(0,1). We compute how well the model estimates the true read counts (Supplemental Figure 8).

For experiments conducted over the defined library, we use the null library to construct a baseline model where the read count in each round follows its own Poisson distribution. The lambda parameter for each distribution was estimated by average read counts (with added pseudocounts for peptides which don’t show up in any round added to make the variance of the distribution in the zeroth round match the mean). When performing MAP estimation, an additional parameter is given to each peptide which indicates whether it comes from this baseline distribution or from the model described above to filter out noise.

RSR values of replicate selections of the defined library are concordant with the first replicate (Pearson and Spearman correlation coefficients 0.81-0.84; Supplemental Figure 9), suggesting selections and RSR determination is reproducible. A subset of sequences is absent from a single replicate due to stochastic dropout, which likely occurs in the initial rounds of selection when each member of the library is present at low frequency.

## Acknowledgments

This work was supported in part by a Schmidt Futures grant to D.K.G. and M.E.B., a Melanoma Research Alliance Grant to M.E.B., and the National Institutes of Health (R01CA218094 to D.K.G. and P30CA14051 to M.E.B.). Siddhartha Jain’s contribution was made prior to him joining Amazon.

## Author contributions

Z.D. provided the analysis of the peptide display data under the supervision of D.K.G., while B.D.H. performed the assays under the supervision of M.E.B. H.Z., B.C., S.J., and B.D.H. performed many initial explorations of the problem and provided preliminary analysis. Z.D., B.D.H., and D.K.G. wrote the manuscript with input from the coauthors.

