## Supplemental Figures 1-9 for "Machine learning optimization of peptides for presentation by class II MHCs"

**Supplemental Figure 1**

**a)**

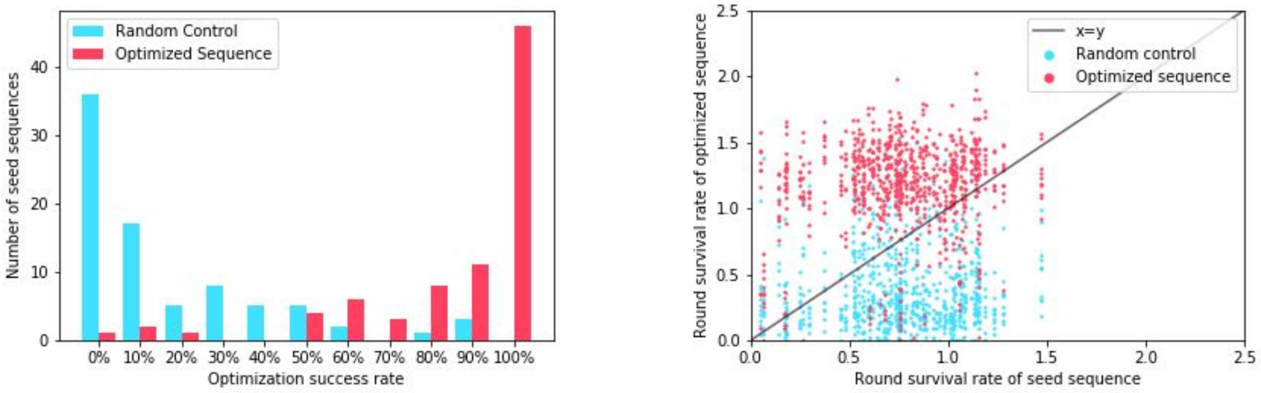


**b)**

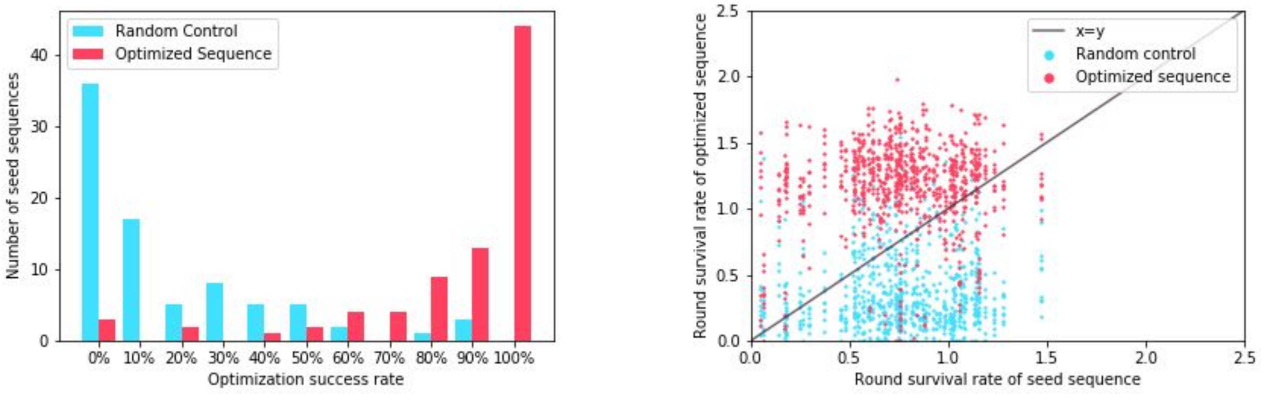


**c)**

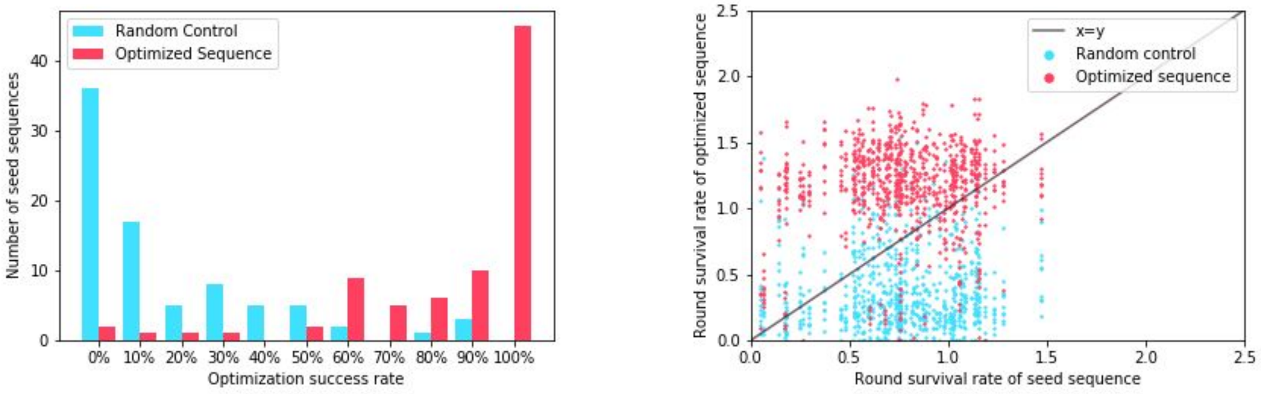


**Number of sequences that exhibit improvement for other optimization methods in DR401.** These depict the same plots as Figure 3, but for different optimization schemes for HLA-DR401. **a)** PE under the categorical model. **b)** UCB under the Gaussian model. **c)** UCB under the categorical model.

**Supplemental Figure 2**

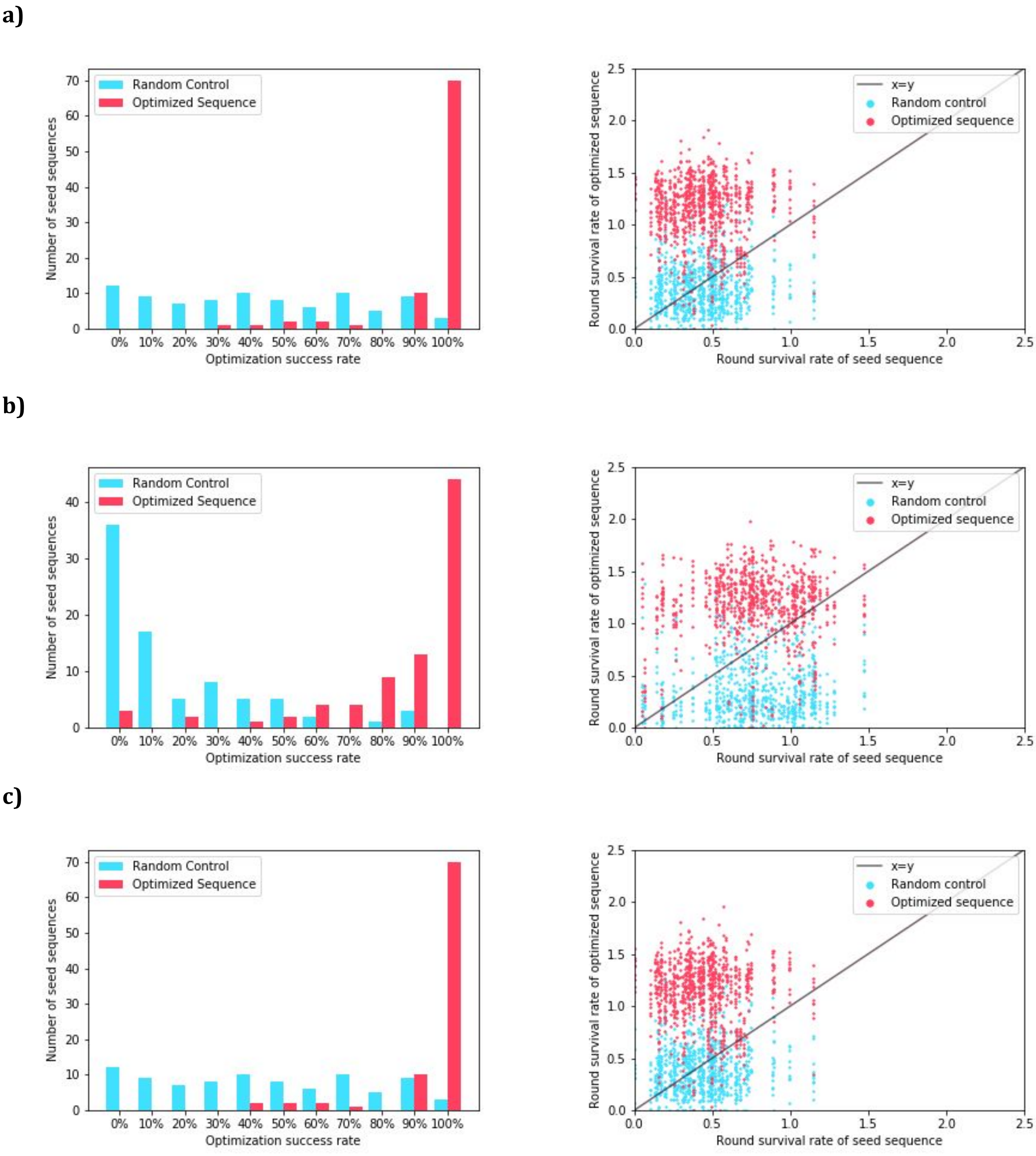


**Number of sequences that exhibit improvement for other optimization methods in DR402.** These depict the same plots as Figure 3, but for different optimization schemes for HLA-DR402. **a)** PE under the categorical model. **b)** UCB under the Gaussian model. **c)** UCB under the categorical model.

**Supplemental Figure 3**

**a)**

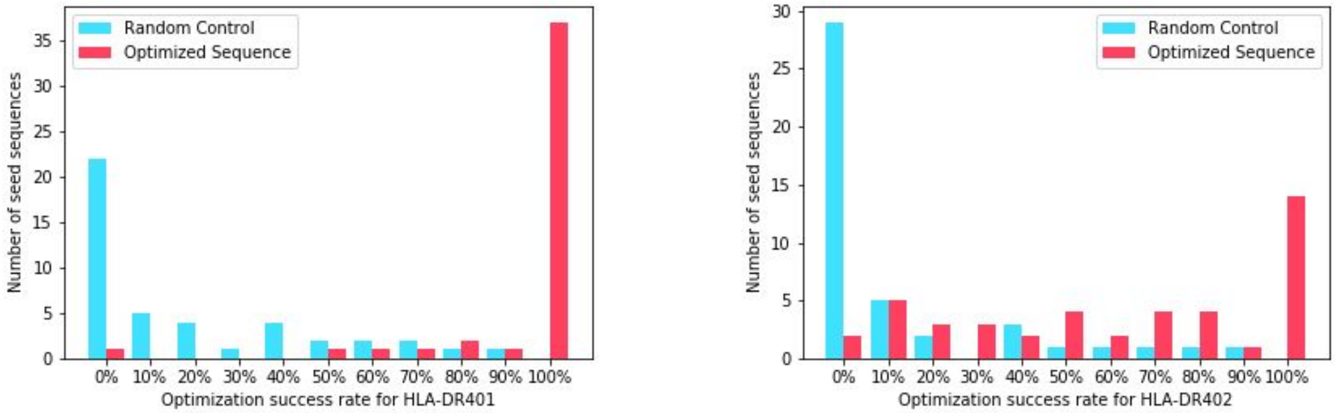


**b)**

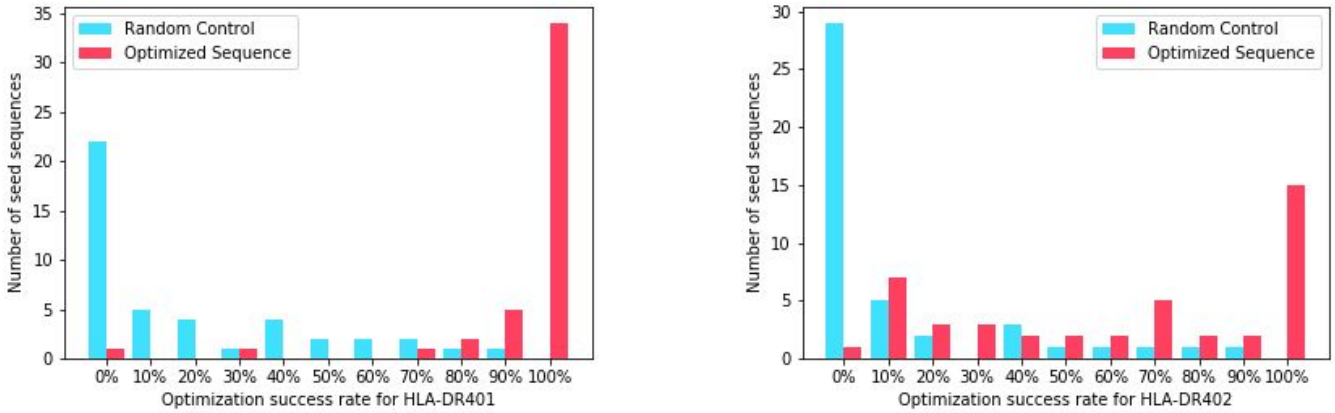


**c)**

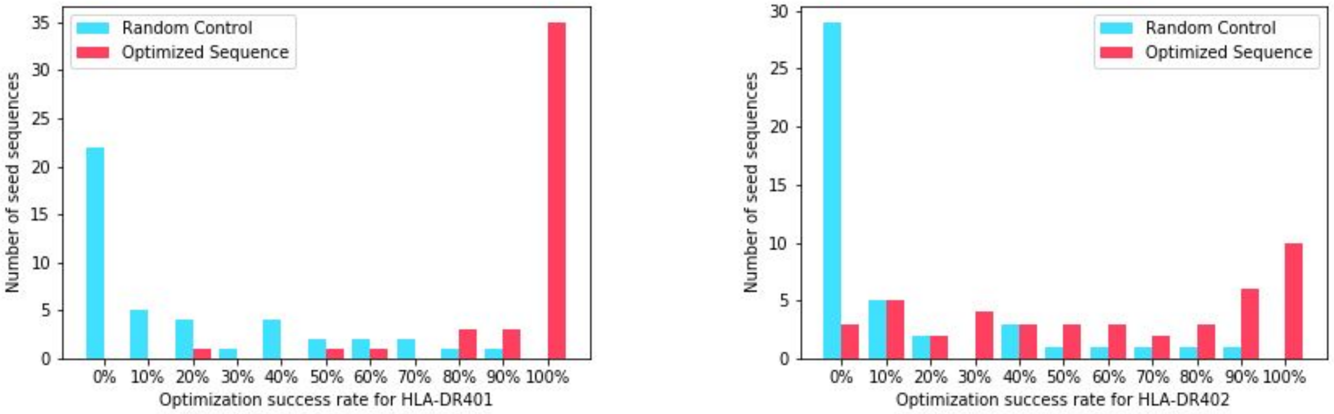


**Multiple allele optimization improvements for other optimization methods.** The same plots as Figure 4a/b, but for different optimization schemes. **a)** PE under the categorical model. **b)** UCB under the Gaussian model. **c)** UCB under the categorical model.

**Supplemental Figure 4**

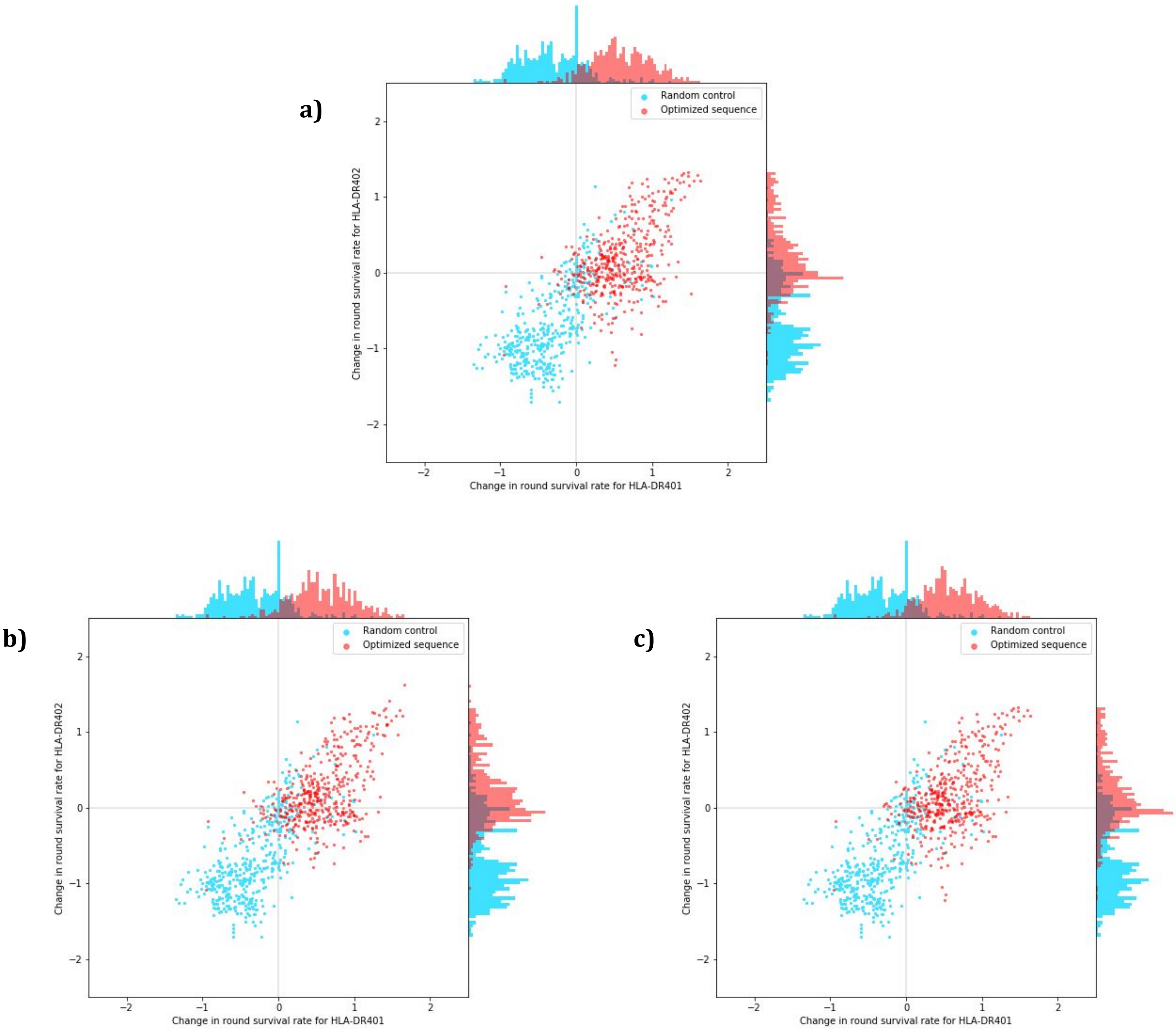


**RSR changes in multiple allele optimization for other optimization methods.** The same plots as Fig 4d, but for different optimization schemes. **a)** PE under the categorical model. **b)** UCB under the Gaussian model. **c)** UCB under the categorical model.

**Supplemental Figure 5**

a)

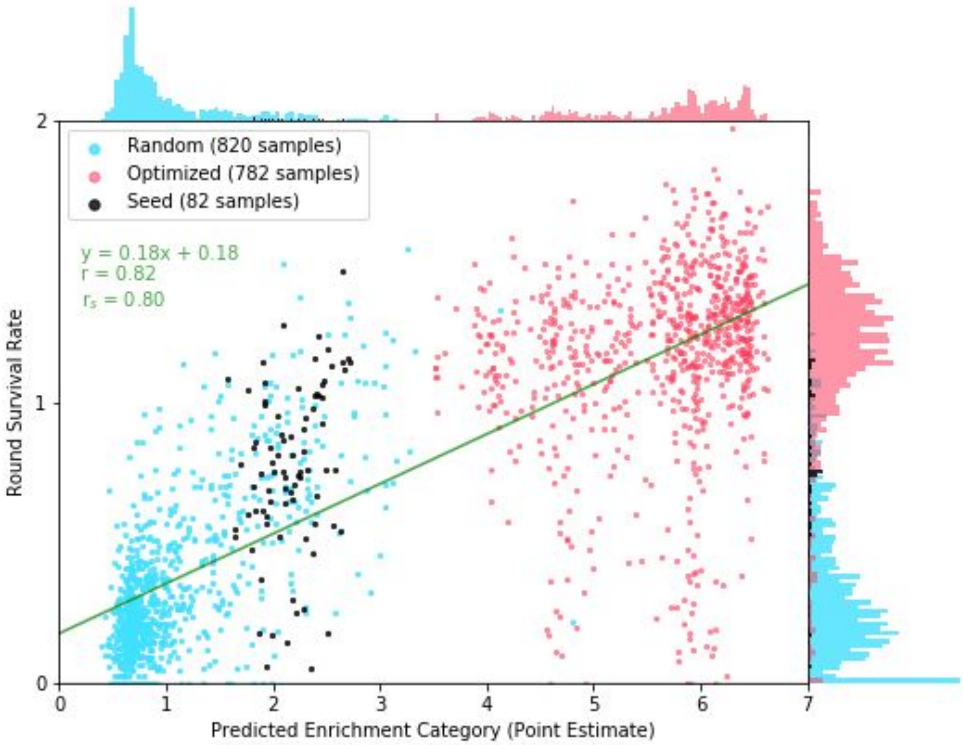


b)

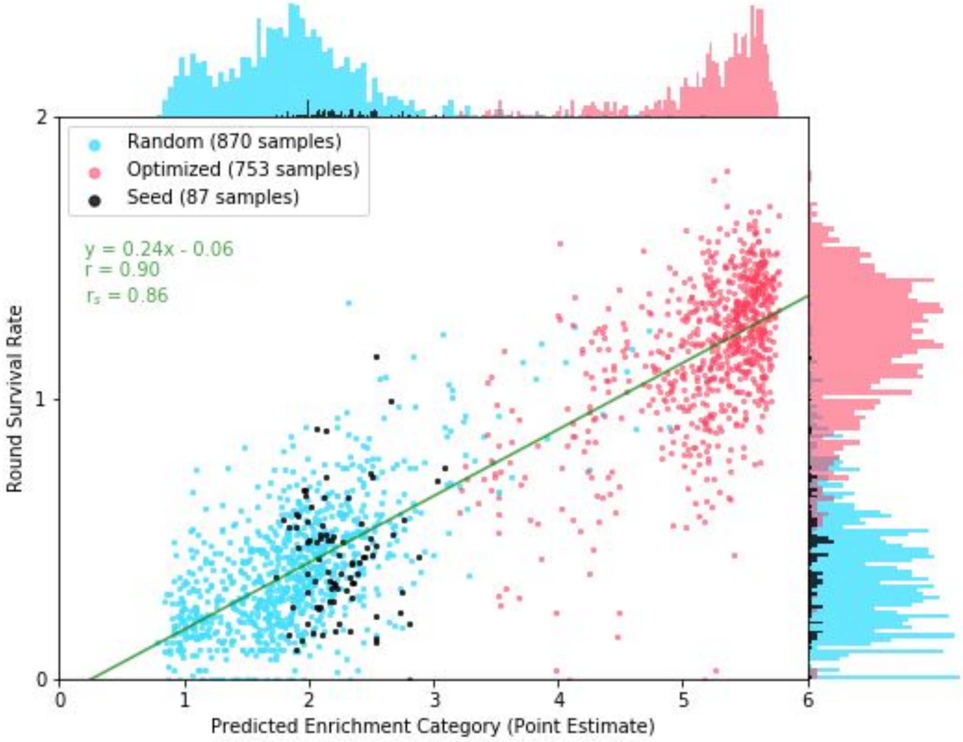


**PUFFIN enrichment predictions correlate strongly with round survival rate. a)** The predicted enrichment values of the categorical model trained for HLA-DR401 is plotted against the measured RSR for HLA-DR401 for seed sequences, sequences optimized using PE with the Gaussian model, and randomly perturbed sequences. The line of best fit obtained from linear regression is shown alongside the Pearson correlation coefficient r and Spearman correlation coefficient $r_s$. **b)** The predicted enrichment values of the categorical model trained for HLA-DR402 is plotted against the measured RSR for HLA-DR402.

**Supplemental Figure 6**

**a)**

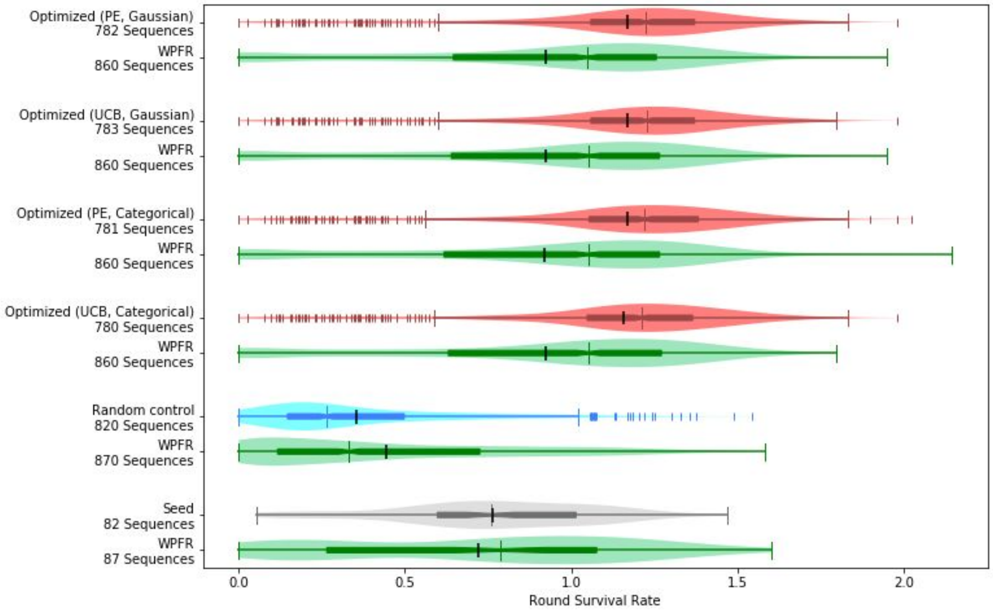


**b)**

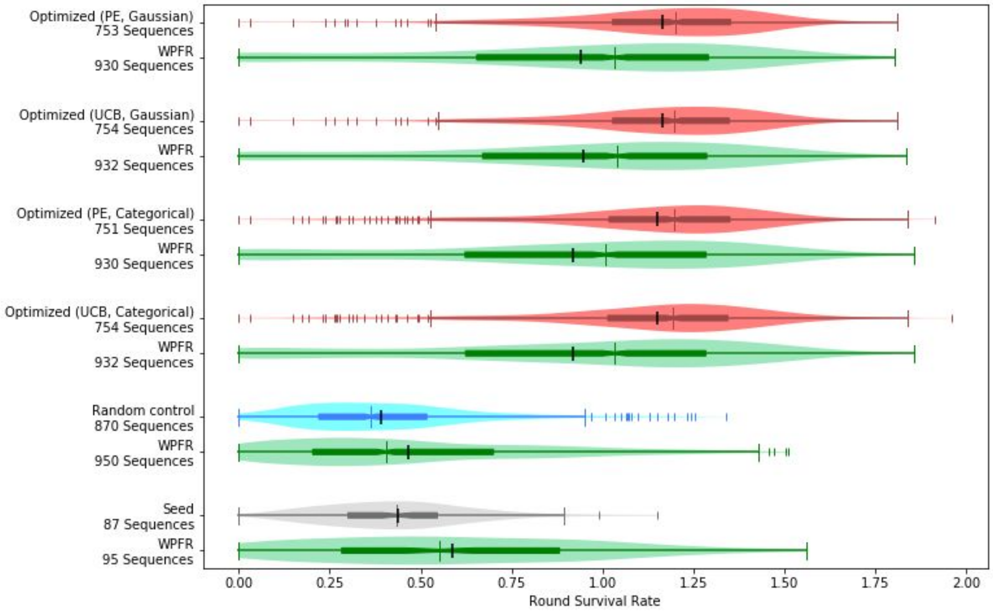


**Comparing round survival rate of various groups with invariant peptide flanking residues and wild type peptide flanking residues.** From top to bottom, the first four pairs of groups depicted in that figure are optimized sequences, and the pairs below them are sequences with random anchor mutations and seed sequences. The upper group of each pair contain IPFR flanked sequences and are the same as those in Figure 2, while the lower group in green contain WPFR flanked sequences. The distributions of RSR for **a)** HLA-DR401 and **b)** HLA-DR402 are plotted for these groups. Each plot is a combination of a box plot and a violin plot, where the distribution is shown by the violin plot in a lighter color, and the box plot shows the middle quartiles in a darker color along with the median. The mean is indicated by a black vertical line. Flier points are marked with the "|" symbol.

**Supplemental Figure 7**

**a)**

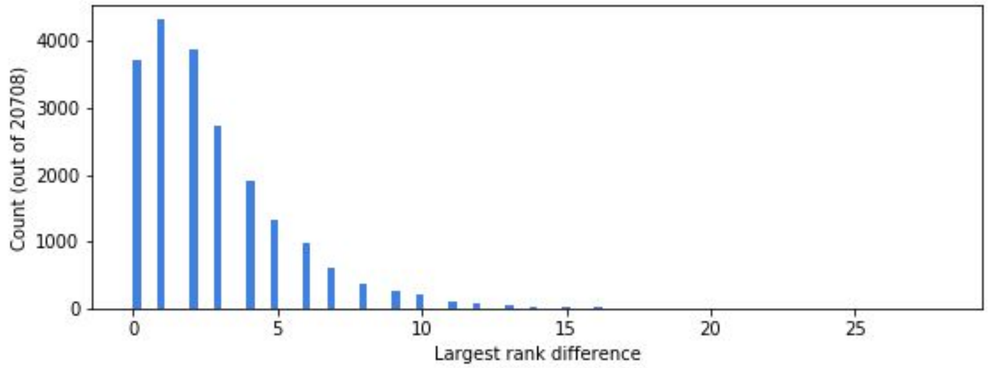


**b)**

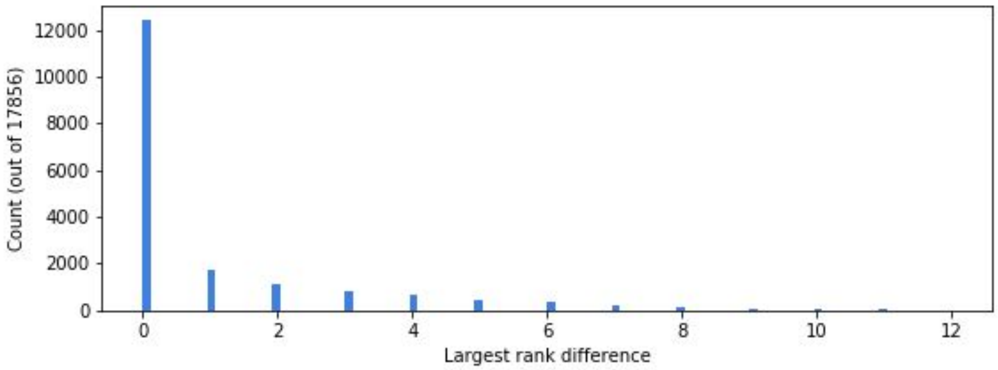


**c)**

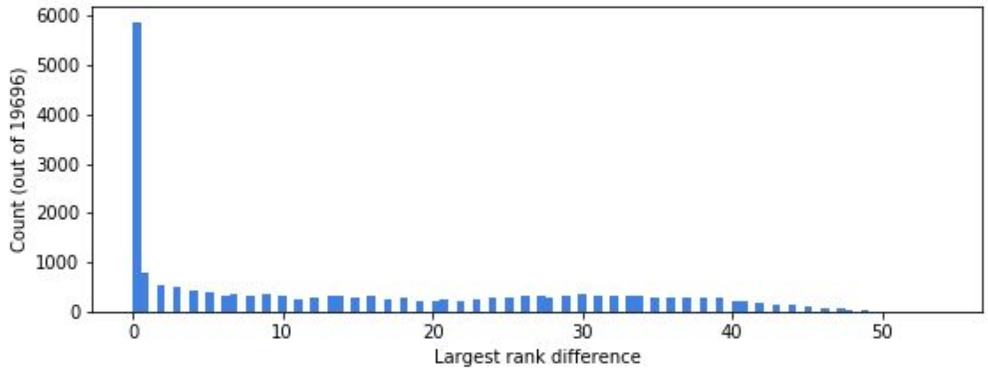


**d)**

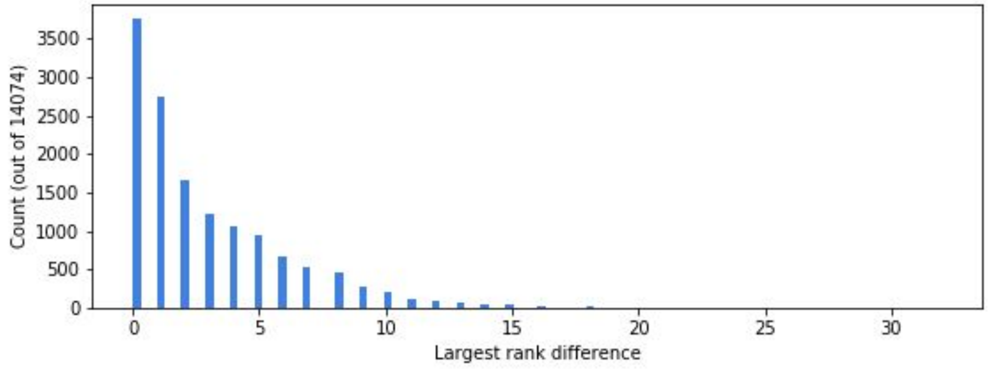


**RSR convergence.** We estimated RSR values for each set of data 23 times. For each run, we obtain an assignment from a read count vector to an RSR value. This gives us an ordering over the read count vectors, and allows us to assign a rank to each unique read count vector. For each read count vector, we compute the largest discrepancy in its rank between the 23 rounds. The histogram depicts the distribution of these discrepancies. **a)** Validation sequences for HLA-DR401. **b)** Validation sequences for HLA-DR402. **c)** Training sequences for HLA-DR401. **d)** Training sequences for HLA-DR402.

**Supplemental Figure 8**

**a)**

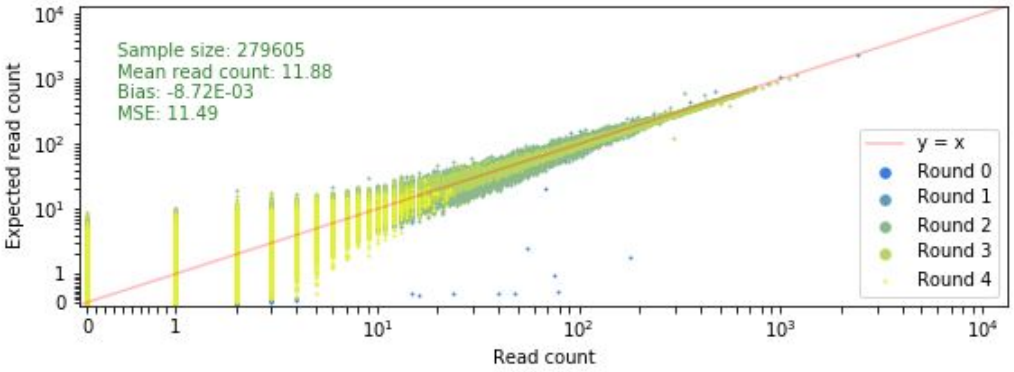


**b)**

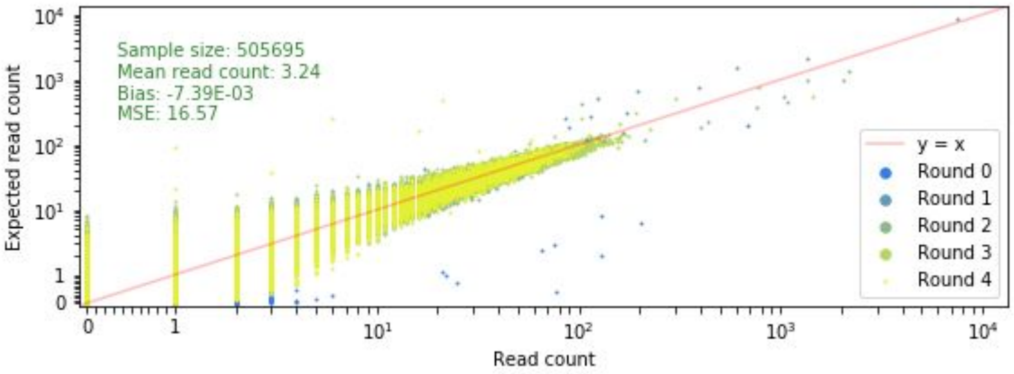


**c)**

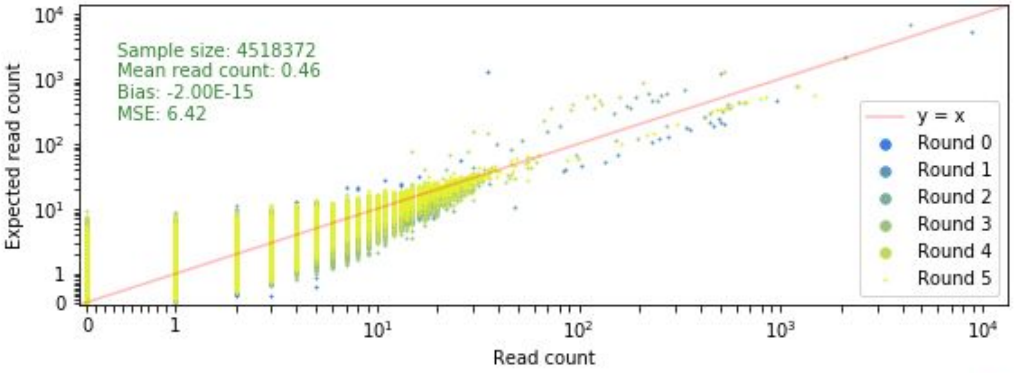


**d)**

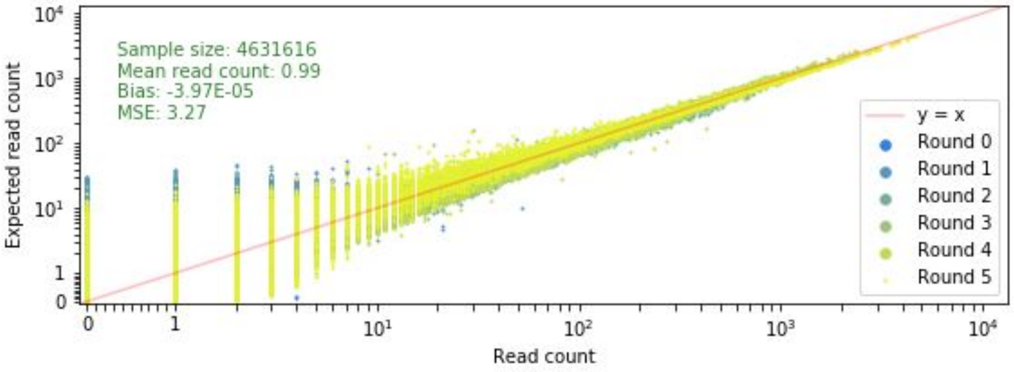


**RSR model fit.** Given the estimated parameters of our MAP estimate, we plot actual read count against the expected read count under the model. **a)** Validation sequences for HLA-DR401. **b)** Validation sequences for HLA-DR402. **c)** Training sequences for HLA-DR401. **d)** Training sequences for HLA-DR402.

**Supplemental Figure 9**

**a)**

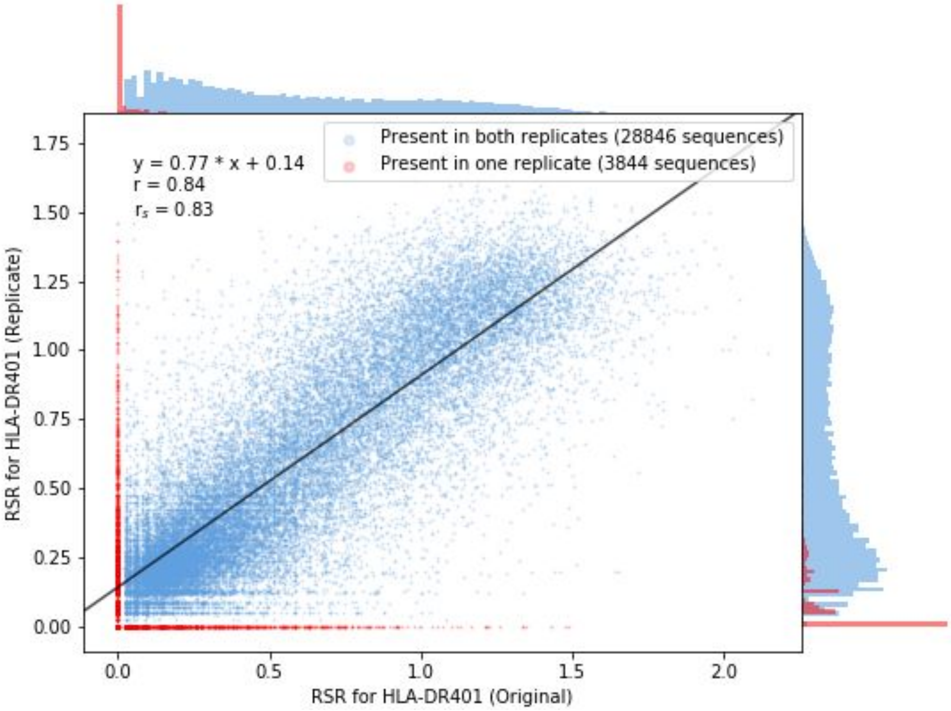


**b)**

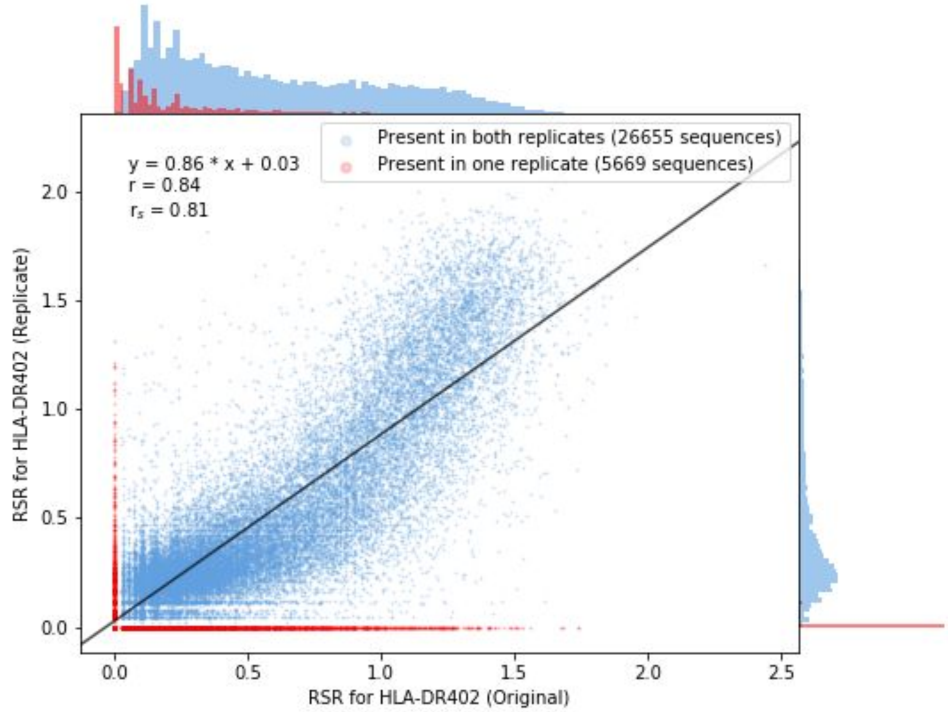


**Validation through replicates.** We validated the RSR values from the validation round by performing a second replicate. **a)** We plot the RSR values between the original and replicate rounds for HLA-DR401. If a point is not present in one of the experiments, it is given a value of 0 and marked in red. The line of best fit obtained from linear regression for points that were present in both experiments, and is shown alongside the Pearson correlation coefficient r and the Spearman correlation coefficient $r_s$. **b)** We plot the RSR values between the original and replicate rounds for HLA-DR402.
